# The ABC transporter MsbA in a dozen environments

**DOI:** 10.1101/2024.06.20.599867

**Authors:** Lea Hoffmann, Anika Baier, Lara Jorde, Michael Kamel, Jan-Hannes Schäfer, Kilian Schnelle, Alischa Scholz, Dmitry Shvarev, Jaslyn E. M. M. Wong, Kristian Parey, Dovile Januliene, Arne Moeller

**Affiliations:** Osnabrück University, Department of Biology/Chemistry, Structural Biology Section, 49076, Osnabrück, Germany; Center of Cellular Nanoanalytics Osnabrück (CellNanOs), 49076 Osnabrück, Germany

## Abstract

High-resolution structure determination of membrane proteins typically requires isolation from the native lipid bilayer and reconstitution into artificial membrane mimics. For this purpose, numerous detergents, amphipols, polymers and membrane scaffold proteins are available. The choice of the specific membrane substitute can strongly affect the protein’s specific activity, stability and conformational spectrum, potentially leading to errors or misinterpretation during analysis. The bacterial ATP-binding cassette transporter MsbA is a prominent example of such environment-specific bias, resulting in apparent conformational and activity responses.

Here, we present a systematic analysis of the conformational spectrum of MsbA, stabilized in a dozen environments, using cryo-EM. Our data show pronounced structural feedback of the ABC transporter to the respective membrane mimetics. Detergents generally favour a conformation with wide separation of the nucleotide-binding domains, while nanodiscs induce the narrow conformation. Notably, only three of the dozen tested environments allow MsbA to sample the functional conformational spectrum, enabling full movement of the nucleotide-binding domains between narrow and wide inward-facing conformations. We expect this study to serve as a blueprint for other membrane proteins, even where the structural reaction to the hydrophobic environment is not directly visible but still critical for the proteins’ function.

## Introduction

Fueled by ATP binding, hydrolysis, and the release of the hydrolysis products, ATP-binding cassette (ABC) transporters undergo significant conformational changes between inward-(IF) and outward-facing (OF) conformations to transport substrates through the lipid bilayer ^1^. Structural studies are key to unravelling the mechanistic details of the transport mechanism, as highlighted by the recent avalanche of ABC transporter cryo-EM structures in different conformations and nucleotide-bound states ^1,2^. Fundamental to cryo-EM single particle analysis is the purification of the target protein from the plasma membrane and stabilization of the isolated protein particles in artificial membrane mimics ^3^. Numerous such substitutes are currently available and used for structure determination ^3–5^. For cryo-EM studies of ABC transporters, the most popular are conventional nanodiscs based on membrane scaffold proteins (MSP) ^6^ and the detergent digitonin (including its synthetic substitute glycol-diosgenin (GDN))^2^. Maltosides, like Lauryl Maltose Neopentyl Glycol (LMNG) and n-Dodecyl-beta-Maltoside (DDM), are also commonly used and significantly cheaper. Alternative mimics, such as amphipols, saposin-, or peptide-based nanodiscs (beta-peptides and peptidisc), are less common ^7–9^.

For all nanodiscs, the most obvious advantage over detergents is the ability to modify the composition of the lipids and tailor them to the requirements of the target protein. Furthermore, once reconstituted, the proteins can be treated as soluble targets in detergent-free solvents, making the proteins amenable to, for example, mass photometry ^10^ and allowing for lower concentrations in cryo-EM sample preparation. Nevertheless, conventional nanodiscs depend on a detergent-based solubilization step, which challenges the co-purification of native lipids. Styrene maleic acid co-polymer lipid particles (SMALPs) ^11^, amphipols ^12^ and Salipro-based DirectMX technology ^13^ offer additional solutions, allowing the direct extraction of the protein along with native lipids from the membrane. Still, their application has been limited to a few successful cases ^14^.

Notably, the activity of an ABC transporter is typically not only driven by the presence of ATP and a transport substrate but also by their specific lipidic environment ^7,15–17^. For example, the ATPase activities of nanodisc-embedded ABC transporters are significantly higher compared to detergent. However, as activities in nanodiscs even exceed those measured in liposomes ^16^, this effect cannot be accounted for only by the presence of lipids, but additional factors must also influence the membrane protein.

The ABC transporter MsbA exhibits a pronounced phenotype when reconstituted into different membrane mimics, rendering it an ideal example to study environment-induced effects ^17–19^. In gram-negative bacteria, MsbA flops the rough lipopolysaccharide (LPS) through the inner membrane to the periplasm, where O-antigen units are attached. The resulting smooth LPS is further transported to the outer membrane to create a protective layer and to reduce the permeability of antibiotics.

While many structures of MsbA have been published at high resolution, its conformational spectrum has been extensively debated. As such, structures deduced from crystals of three orthologous transporters under different nucleotide-bound states revealed marked conformational variances ^20^. Here, an IF_narrow_ structure resembled a conformation typically described for ABC transporters. However, the second IF conformation exhibited nucleotide-binding domains (NBDs) separated by about 70 Å. This unusually wide conformation raised the question of whether it might have been artificially influenced by crystal packing, especially as EPR studies in liposomes only supported distances of about 30 Å ^21^. In contrast, an EPR study on the related ABC transporter LmrA reported a conformational spectrum that would resemble the crystallized states, including wide separations of NBDs ^22^. Later, negative stain EM showed the wide-open conformation of MsbA ^19^, excluding crystallization artefacts as the cause of the large NBD separation. Cryo-EM analyses revealed that the hydrophobic environment, lipid nanodiscs vs. detergent, directly impacts the NBD separation in the IF conformation ^17^. In this study, MsbA exhibited a wide opening only when stabilized in detergent but not in lipid nanodiscs, suggesting that the narrow conformation represents a more physiological state. Similarly, only the narrow IF conformation was observed in peptidiscs ^23^. However, it does not appear that simple, as results obtained by smFRET revealed a wide opening of MsbA in liposomes (Liu et al., 2018) while supporting the finding that this state does not occur in nanodiscs. Adding to the complexity, cryo-EM structures of MsbA show co-purified lipid A sandwiched between the transmembrane domains (TMDs) ^17,18,24^, in the narrow IF conformation but not in the wide open one ^25^. Recently, the application of in-cell DEER measurements, accompanied by careful calibration through cryo-EM, enabled the detection of the wide-open IF conformation inside the cells, ultimately proving its biological relevance ^18^.

The strong phenotype of MsbA, with its eminent reaction to the hydrophobic environment, renders it an ideal target to study the structural response from a membrane protein to various commonly used membrane substitutes and, therefore, poised us to perform a systematic analysis. Here, we present 20 cryo-EM structures of MsbA, determined in 12 different environments. In our data, all detergents show the wide-open conformation with a correlation between critical micelle concatenation (cmc) and the degree of NBD separation. Only LMNG and GDN additionally exhibit a narrow IF-conformation. All nanodisc stabilized MsbA preparations exhibit the narrow conformation exclusively, except for the very large MSP2N2 nanodiscs, which also allow for the wide-open conformation. For all the MSP-based nanodiscs, ATPase activities are comparable but generally much higher than those of detergent systems. In summary, our systematic analysis reveals the structural and functional bias of various membrane mimics, which must be considered for structure determination and interpretation of membrane proteins.

## Results

Here, we used cryo-EM single particle analysis to systematically compare the conformational spectrum of the ABC transporter MsbA in a nucleotide-free state in twelve different environments. For direct extraction of MsbA, *Escherichia coli* membranes were solubilized individually with each tested detergent and subjected to nickel affinity purification (see Methods). The UDM-solubilized sample was further used for peptidisc reconstitution, while the DDM-solubilized sample was used for MSP-based nanodiscs and amphipol A8-35 preparations. Detergent and amphipol 18 solubilized samples were concentrated to 10-14 mg/ml for cryo-EM specimen preparation, while nanodisc, peptidisc and A8-35 reconstituted samples only required 1.8-2.5 mg/ml. After vitrification, the proteins were imaged and processed in a highly standardized manner using cryoSPARC ^26^. The processing workflow was adapted for the final refinements to fit each dataset’s composition and challenges (Fig. S6, S7).

Environments enabling direct extraction from the membrane, such as detergents and amphipol 18, generally favoured the IF_wide_ conformation (Fig. 1-c,f). In contrast, for most nanodisc reconstituted samples, IF_narrow_ was found exclusively (Fig. 2 a-d). Exceptions are the large MSP2N2 nanodiscs in which 30% of MsbA particles displayed the IF_wide_ conformation (Fig. 2 d). The detergents LMNG and GDN contained both IF_narrow_ and IF_wide_ conformations with even distribution (Fig. 1 d,e).

**Fig. 1.**
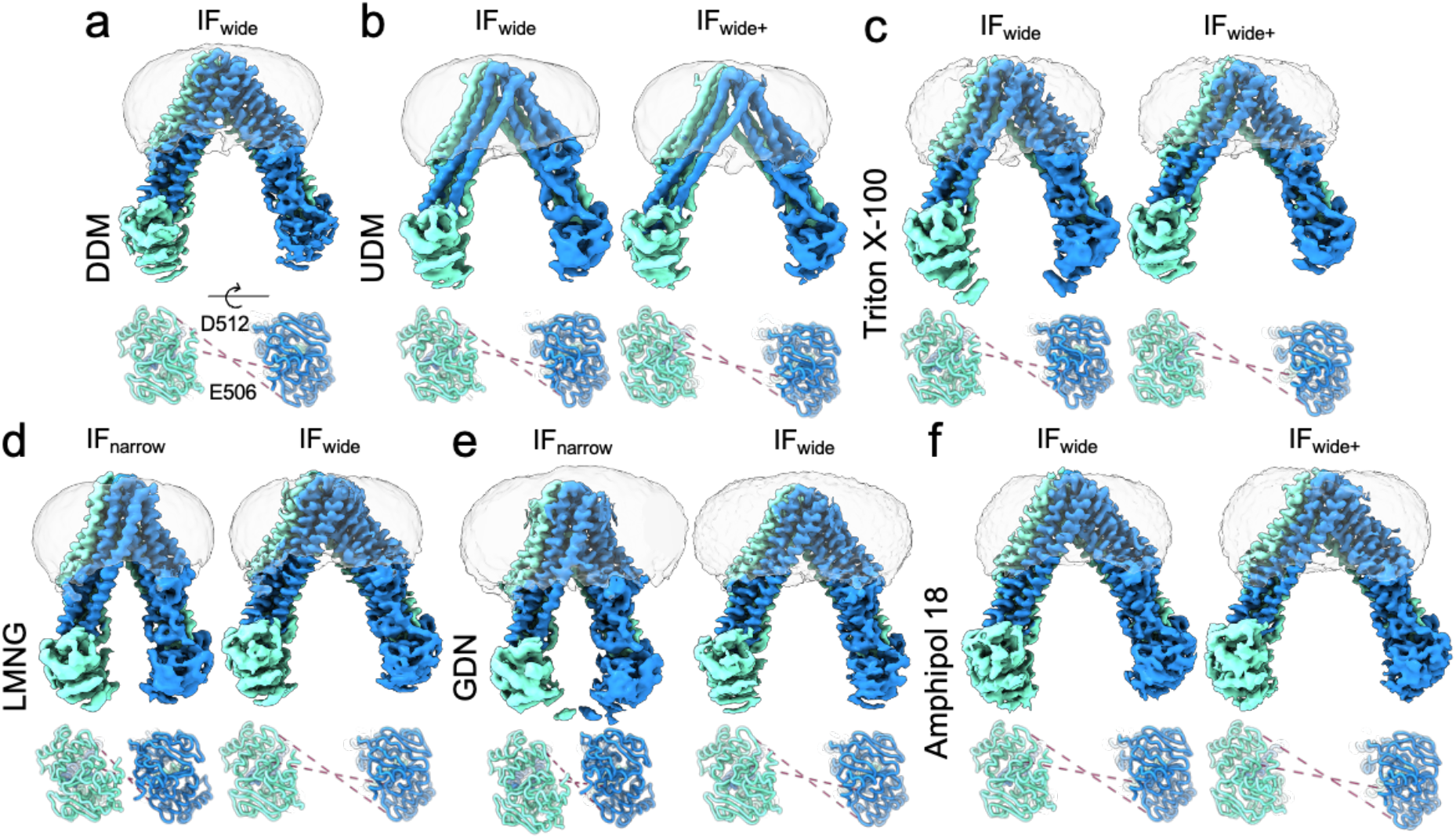
Conformational variety of nucleotide-free MsbA from direct membrane extractions. Cryo-EM densities of MsbA in DDM **(a),** UDM **(b),** Triton X-100 **(c)** and amphipol 18 **(f)** depicting a range of wide IF conformations, GDN **(e)** and LMNG **(d)** exhibiting both IF_narrow_ and IF_wide_ states. The hydrophobic environment is depicted in a light grey transparent surface. The measured distances between residues D512 and E506 are indicated as dashed lines. MsbA densities are coloured by chain ID (cyan and blue).

**Fig. 2.**
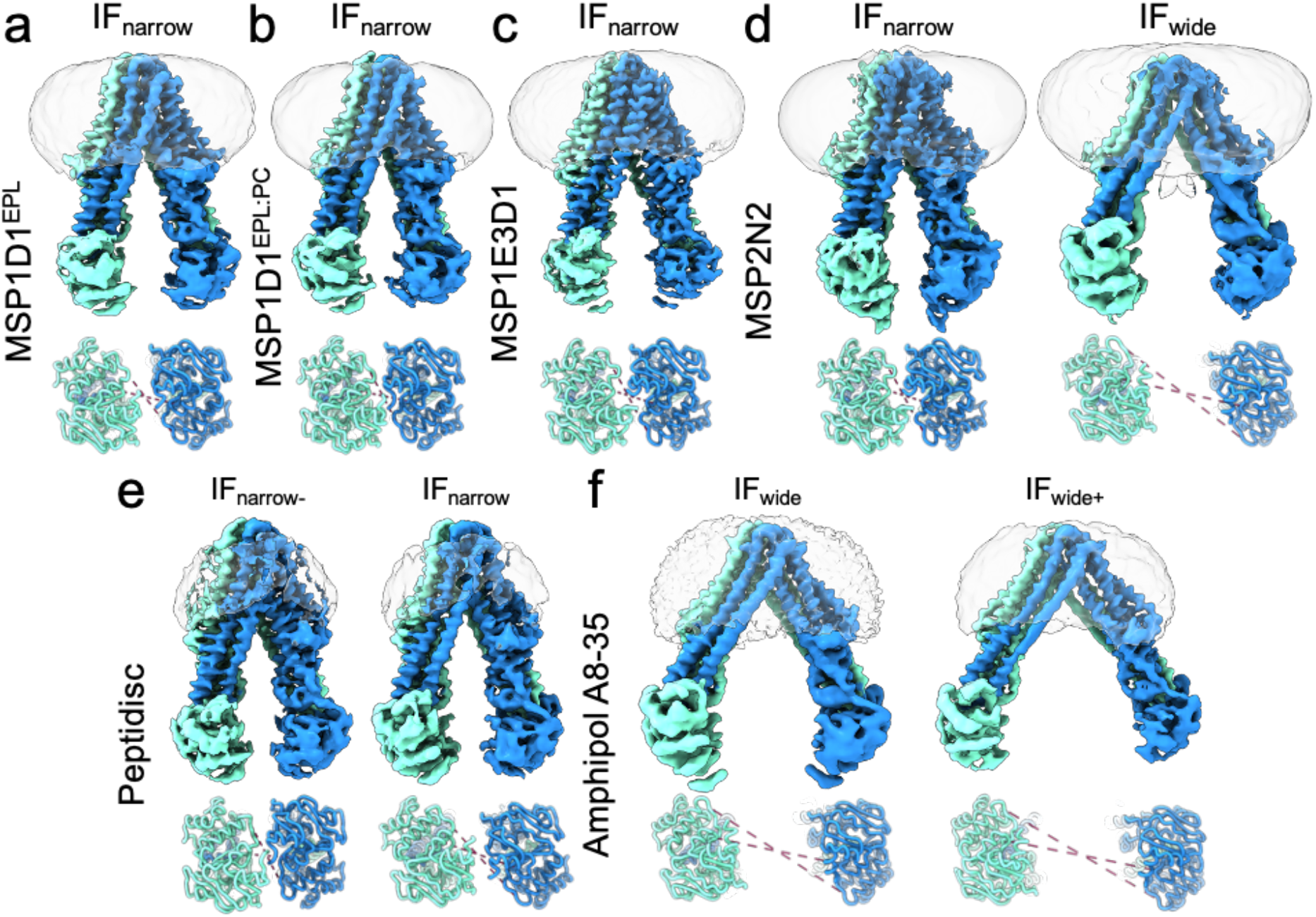
Conformations of MsbA, reconstituted in various membrane mimetics. The density maps of MsbA in the lipid nanodiscs MSP1D1^EPL^ **(a),** MSP1D1^EPL:PC^ **(b)** and MSP1E3D1 **(c),** as well as in the peptidisc **(e)** exhibit only IF_narrow_ conformations. Only in MSP2N2 nanodiscs **(d)** the functional IF spectrum of MsbA is present. Detergent reconstitution to amphipol A8-35 results in IF_wide_ conformations exclusively **(f).** The measured distances between the residues D512 and E506 are indicated as dashed lines. MsbA densities are coloured by chain ID (cyan and blue).

As reported previously, the opening of the transporter is not linear but follows a simultaneous twist and turn motion ^20^. Therefore, to classify the various observed degrees of NBD separation, we measured two symmetric amino acid pairs (E506 and D512 (Fig. 1, 2, Supp Table 2)). All conformations described here likely represent the global average of a dynamic equilibrium of many similar conformations rather than one stable state. This motion is supported by the comparably modest resolution obtained from IF conformations (3.2-4.0 Å vs. 2.6-2.9 Å for OF conformations) ^18,27^.

In detergent, we consistently observed the presence of a wide-open IF conformation with NBD separation ranging from 58 Å to 78 Å. As noted above, only LMNG and GDN solubilized MsbA additionally exhibited an IF_narrow_ conformation with 30 Å to 40 Å separation between the NBDs (Fig. 3). In line with the previously published structures, the inter-NBD distance narrows in nanodiscs and peptidiscs ^17,28^. Alteration of the lipid composition does not seem to affect the opening degree (Fig. 2 a,b). However, the size of the belt protein influences the conformational spectrum. While MSP1D1 and MSP1E3D1 nanodiscs (diameter – 9.5 and 13 nm respectively) ^29^ restrict MsbA to an IF_narrow_ conformation (Fig. 2 a-c), the larger MSP2N2 nanodiscs (diameter – 17 nm) ^30^ allow for an IF_wide_ conformation (Fig. 2 d) with 60 Å NBD separation.

**Fig. 3.**
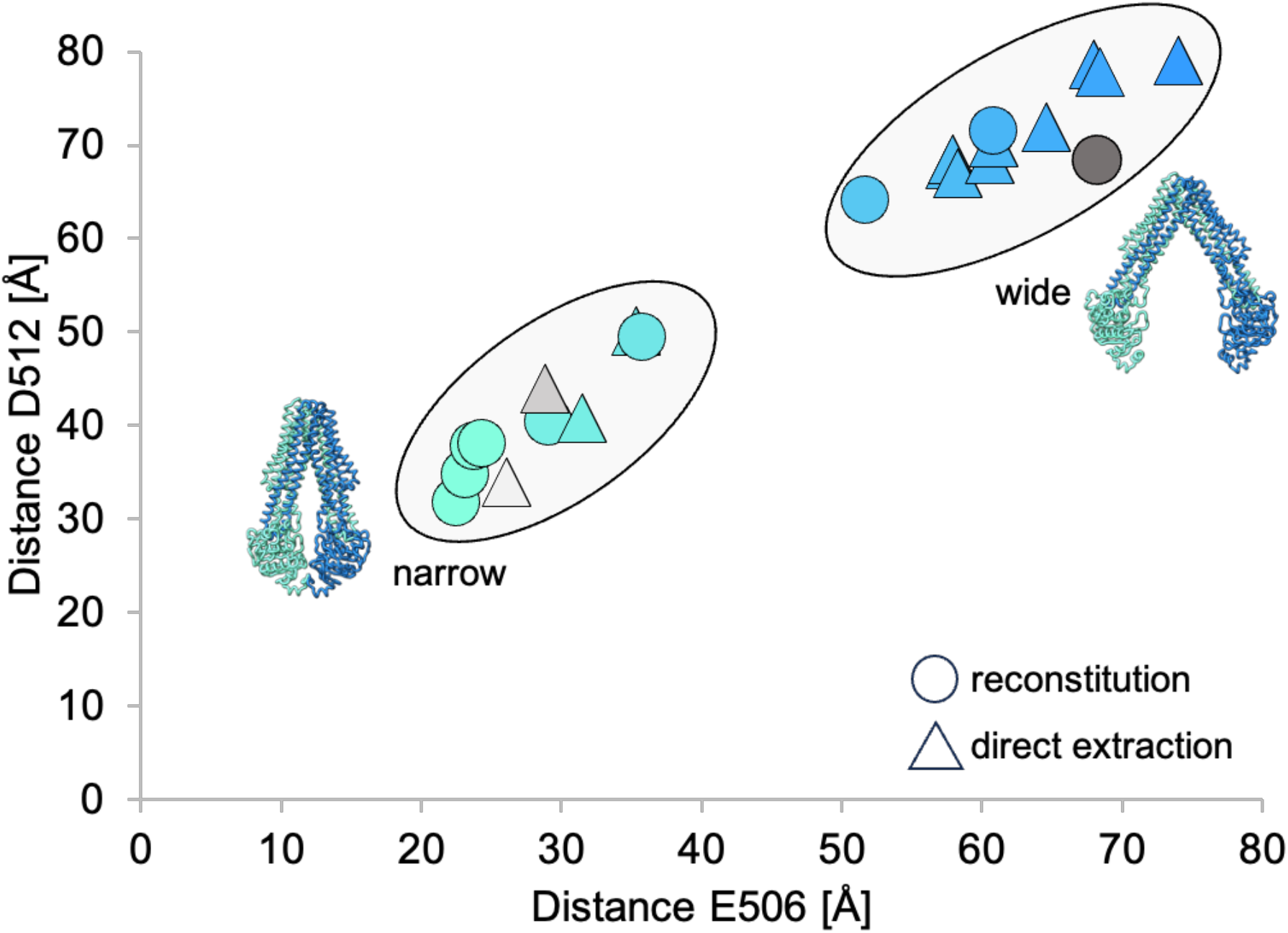
MsbA conformation clusters. Distance measurements in the NBDs between MsbA chains A and B at residues D512 and E506 are plotted against each other. Structures of MsbA in cartoon representation exemplify narrow and wide conformations. Triangles indicate direct protein extractions and circles – reconstitutions. Previously published structures are displayed: light grey – MsbA in nanodisc (5TV4), grey – MsbA in peptidisc (6UZ2) and dark grey – MsbA in DDM (8DMO) (cf. Sup. Table 2).

In our structures, we generally do not observe a significant increase in bound lipids when comparing the highest-resolution structures of nanodisc-embedded and detergent-stabilized MsbA. As previously shown for the pentameric ligand-gated ion channel pLGIC ^31^, MsbA interacts directly with the belt proteins like MSPs or through protein-lipid-protein interaction. Therefore, the protein is not freely floating within the nanodisc but preferentially contacts the amphiphilic boundary of the discs (Fig S4).

Despite the different strategies used for amphipol samples, either reconstitution into A8-35 from detergent (Fig. 2 f) or direct extraction from the membrane with amphipol 18 (Fig. 1 f), the observed conformations were similar and comparable to the Triton X-100-purified sample, displaying the largest NBD separation (73-76 Å) of all samples tested.

While the wide-open conformation dominates in UDM and amphipol, we also observe a small, narrow population (Fig. S1). However, we must exclude this from our statistics due to the small particle counts and low resolution.

We do not observe a clear correlation between the activity and conformation. As previously published, the ATPase activity is generally much higher in nanodiscs than detergents ^7^. Also, the highest activity of MsbA coincides with the reconstitution in the tightest MSP1D1 nanodiscs. In contrast, the lowest activity was observed in amphipols with large NBD separation (Fig. 4). In all detergent-solubilized samples, the activity was similar regardless of the conformations detected by cryo-EM. An exception was the UDM sample, which had an unexpectedly low activity comparable to amphipols. The ATPase activity in the peptidisc was comparable to detergents. Of note, we observed precipitation in the peptidisc-reconstituted sample after adding Mg^2+^, which could have decreased the protein concentration in the solution and, thereby, negatively affected the accuracy of the calculation.

**Fig. 4.**
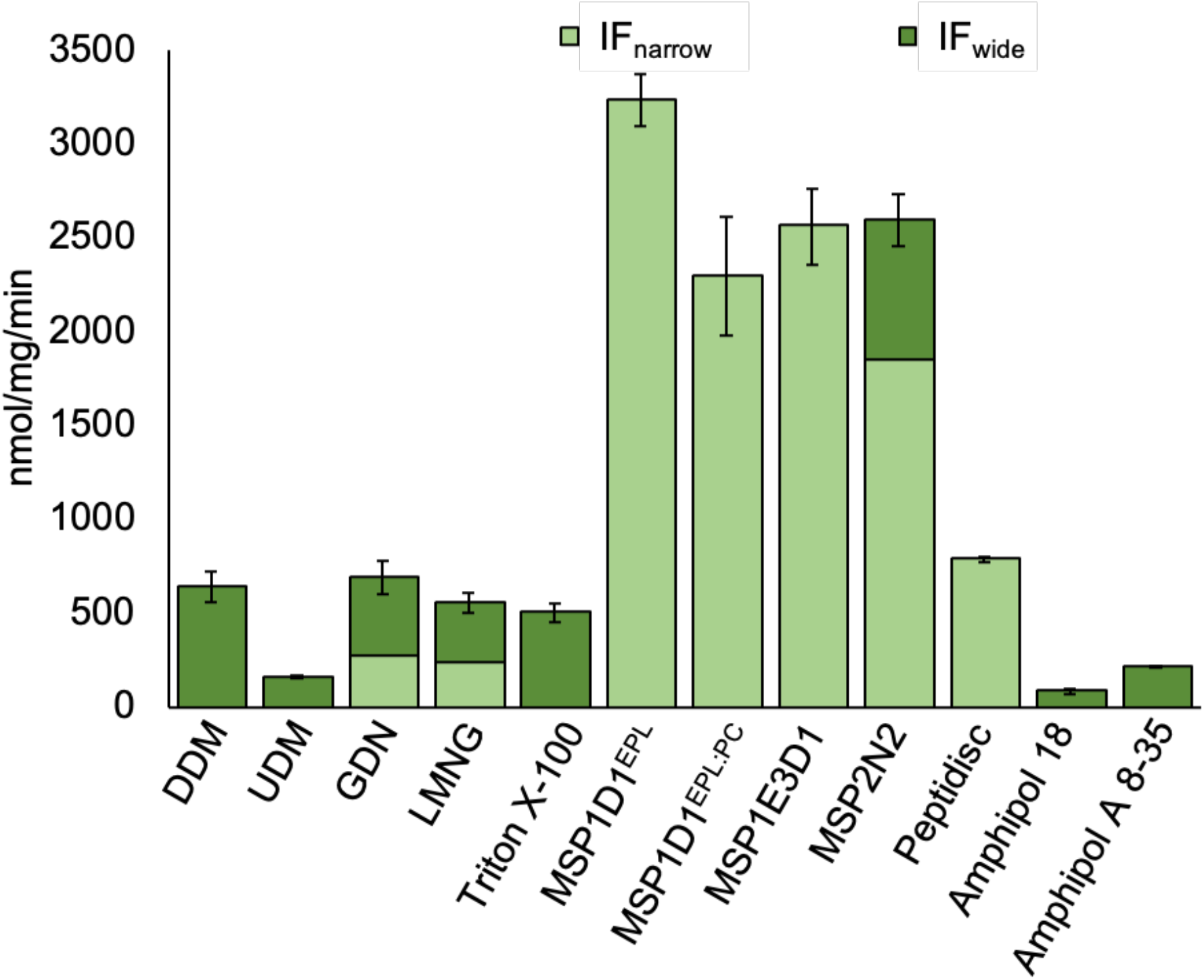
ATPase activity measurements and relative conformations. The ATPase activity was measured using a molybdate-based colourimetric assay and presented as the amount of phosphate released per mg of MsbA per min (mean ± standard deviation; n = 3). The bars are partitioned by the percentage of the conformations present in the dataset and do not represent the activity for each conformation.

Moreover, the peptidisc samples were reconstituted from the low-activity UDM preparations, potentially accounting for the 3-fold lower activity than the MSP-based nanodisc preparations reconstituted from DDM.

The bound LPS precursor is present in all IF_narrow_ conformations (Fig. S2), irrespective of the hydrophobic environment. Such a density is far less evident for IF_wide_, only appearing at a low-density threshold, thus insufficient for model building and quantitative assignment.

## Discussion

Several reports have recently highlighted significant structural responses of membrane proteins to various hydrophobic environments. As such, the transmembrane helices of the acetylcholine receptor exhibit different poses before detergent extraction or after and within a nanodisc ^32^. Similarly, the pentameric ligand-gated ion channel pLGIC exhibits distinct conformations in various nanodiscs ^31^. MsbA is another example with a pronounced structural response to various hydrophobic environments ^16–18^. Here, we investigated the conformational reaction of *E. coli* MsbA to twelve different membrane mimetics and compared it to the respective ATPase activity. The IF conformation was previously reported to be narrow for nanodisc-based reconstitutions and wide for detergent counterparts. In-cell DEER measurements have recently revealed that the entire conformational spectrum of MsbA is present inside the *E. coli* membrane, with the wide open conformation being dominant ^18^. Accordingly, the MsbA system, with its apparent phenotype, provides a unique opportunity to analyze the specific structural response to various hydrophobic environments systematically.

Our survey shows that all nanodisc-based samples result in narrow IF conformations, and only the largest tested nanodisc, MSP2N2, additionally exhibits the IF_wide_ conformation for 30 % of particles (Fig. 2, 4). Contrarily, detergents strongly favour a wide IF conformation apart from LMNG and GDN, which displayed an equal distribution between IF_narrow_ and IF_wide_ (Fig. 1, 4). Interestingly, we observed a correlation between the critical micelle concentration (CMC) and the degree of NBD separation: the higher the CMC of the respective detergent – the larger the separation of the NBDs.

While examining the intracellular gate of the IF_wide_ and IF_narrow_ structures, only the narrow state contains clear density for an LPS-precursor sandwiched between the TMDs (Fig. S2). In turn, tightening of the ligand binding site stabilizes a binding pose, which increases the local resolution and resolvability of the bound state.

The IF_wide_ conformations are generally more flexible and display more variation in their opening. A consequence of the wide NBD separation is an enlargement of the binding pocket, which may tolerate multiple binding modes of the rough LPS or incomplete occupancy, resulting in loss of signal during averaging and challenging unambiguous LPS assignments. Additionally, treatment with harsher detergents could remove the otherwise tightly bound transport substrate. Lipid-like compounds, such as LPS, might be especially prone to detergent-induced diffusion from its binding site. As the LPS precursor was consistently detected in the nanodisc reconstructions, we can exclude that at least UDM/DDM caused LPS removal. However, the extensive NBD separation and absence of apparent ligand density observed in amphipol 18 and Triton X-100 reconstructions suggest that LPS removal may have occurred in these instances.

All systems tested here are influenced by a detergent (or amphipol)–based solubilization step. SMALPS and DIBMALPS have been shown to overcome this limitation ^33,34^. However, they carry additional complexities, such as sensitivity to divalent cations, substantial disc-heterogeneity or low yields. We could not obtain samples of sufficient amounts and quality to perform high-resolution studies of MsbA with these polymer-based membrane mimics. Further optimization is needed to achieve samples compatible with high-resolution cryo-EM reconstructions of ABC transporters ^35^.

According to our data, only LMNG, GDN, and MSP2N2 nanodiscs allow MsbA to adopt a wide range of naturally observed IF conformations. All other environments funnelled the spectrum to IF_narrow_ or IF_wide_ (Fig. 1, 2). Likewise, different hydrophobic environments affect the ATPase activity of MsbA (Fig. 4). As MSP systems favour the narrow conformation, we initially reasoned that the short distance between the nucleotide-binding sites (NBS) amplifies the frequency of dimerization, thus increasing the ATPase activity. However, a comparison of the available datasets reveals that the explanation for the observed increase in activity cannot be that simple. First, the activity of MsbA in MSP2N2 is equally high compared to MSP1E3D1 (Fig. 4), which exhibits only the narrow conformation (Fig. 2 c). Likewise, the activity of the detergents is generally low (Fig. 4), even though LMNG and GDN equally represent narrow and wide conformations (Fig. 1 d, e). Also, the influence of the lipid environment itself cannot serve as the final answer. First, MsbA measured in liposomes exhibits a lower activity than in MSP-based environments ^16^. In PC-supplemented nanodiscs, the activity is also lower than standard *E. coli* polar lipid nanodiscs (Fig. 4), suggesting that the lateral pressure may impact activity more than a specific lipid type. Secondly, adding lipids to the detergent samples did not increase its activity (Fig. S5). Finally, the substrate LPS stimulates ATPase activity ^36,37^. Still, it is also present in IF_narrow_ conformations of LMNG or GDN solubilized particles, which account for about 50% of their total population.

In conclusion, we suggest an additive effect: on the one hand, the short distance between the NBS in the IF-narrow conformations supports fast dimerization. On the other hand, the presence of lipids seems to have an added benefit for activity. We also cannot exclude the inhibitory effect of detergent molecules, as reported for the functionally related human P-gp ^38^. While we cannot provide an ultimate explanation for the observed differences in the ATPase activity, an essential outcome of this comparison is that a high ATPase activity is not a reliable indicator to assure a native conformational spectrum.

The best environment for a specific system may depend on the scientific question. Based on our data, only MSP2N2 nanodiscs provide high MsbA activity and a functional conformational spectrum. This is in line with the recent report on *pLGIC* ^31^ and is to be expected as the belt proteins’ influence on the protein must decrease with growing diameter. However, the increased size and heterogeneity of the nanodisc severely hampers throughput during image processing. The strong signal of the nanodisc compared to the size of the transporter required customized masking to separate the protein signal from the environment, significantly slowing down the processing speed (Fig. S3). Notably, the final results were comparable with the LMNG-stabilized MsbA (Fig. 2 d, 1 d). Therefore, while extraction with LMNG comes at the cost of low activity, it requires significantly fewer movies and particles (Fig. S3), leading to highly efficient processing, which may render LMNG a cost-efficient solution for most cases. In summary, the proper environment choice is sample-dependent, but we hope our analysis might serve as a template for the researcher to choose wisely.

## Acknowledgements

This work was supported by a grant of the DFG (MO2752/3-6), the SFB 1557, the DFG INST190/196-1 FUGG and the RTG 2900 (all A.M.). J.E.M.M.W. received funding through the Novo Nordisk Foundation Grant NNF18OC0030544. J.H.S. is supported by a fellowship from the Friedrich Ebert foundation.

## Author contributions

L.H., D.J. conceived and designed all experiments together with A.M. Cryo-EM analysis and sample preparation was a team effort from all authors. Final samples and datasets were obtained from L.H. (detergent stabilized samples and peptidisc), A.B. (nanodisc stabilized samples), L.J. (amphipol) J.E.M.M.W. (DDM). K.P. built the molecular models. The manuscript was written by L.H., D.J and A.M. with contributions from all authors.

## Competing interests

none

## Methods

### Protein expression and purification using detergents

The pET-19b plasmid, carrying His-tagged MsbA, was a kind gift from Glaubitz group ^39^. It was transformed into competent *E. coli* C41 (DE3) cells, which were grown in LB media with carbenicillin at 37°C till an OD_600_ of 0.6 – 0.8 was reached. The expression of MsbA was then induced with 1 mM IPTG and carried out overnight at 20 °C. The cells were harvested at 6000 g for 10 minutes and resuspended in Lysis Buffer (50 mM Tris/HCl pH 8, 500 mM NaCl, 0.5 mM DTT, 1 mM PMSF, 50 µg/mL DNAse, cOmplete protease inhibitor tablet, 10 % glycerol) and disrupted via sonication. After removing the cell debris by centrifugation (10700 g for 30 min at 4 °C), the supernatant was ultracentrifuged at 140000 g for 1 hour at 4 °C to collect the membrane fraction. The membranes were solubilized overnight at 4 °C while gently stirring in Solubilization Buffer (Lysis Buffer with 10 mM Imidazole and detergent, according to Table 1). The solubilized membranes were ultracentrifuged at 140000 g for 1 hour at 4 °C. The supernatant was mixed with HisPur Ni-NTA Resin and incubated for 2 hours at 4 °C. The resin was applied to a gravity column and washed with Wash Buffer (50 mM Tris/HCl pH 8, 500 mM NaCl, 0.5 mM DTT, 10% Glycerol, 10 mM Imidazole pH 8, detergent according to table 1). His-tagged MsbA was eluted with Elution buffer (Wash Buffer, containing 250 mM Imidazole pH 8). The elution fractions were dialyzed in Dialysis Buffer (20 mM Tris/HCl pH 8, 100 mM NaCl, 0.25 mM DTT, detergent according to table) using a 12-14 kDa MWCO dialysis tube overnight at 4 °C. The dialyzed protein was concentrated with Vivaspin Turbo 4 concentrator (Sartorius) with a 100 kDa MWCO.

Concentrated samples were further purified via size exclusion chromatography using either Superdex 200 Increase 5/150 GL or Superdex 200 Increase 10/300 GL column. Peak fractions were collected and concentrated to the desired concentration.

**Table 1:** Detergent concentrations in protein purification buffers.

| Detergent | Solubilization Buffer Concentration (%) | Wash Buffer Concentration (%) | Elution Buffer Concentration (%) | Dialysis Buffer Concentration (%) |
| --- | --- | --- | --- | --- |
| DDM | 1.25 | 0.1 | 0.1 | 0.05 |
| UDM | 1.25 | 0.3 | 0.3 | 0.15 |
| LMNG | 1 | 0.001 | 0.001 | 0.003 |
| GDN | 1 | 0.004 | 0.004 | 0.004 |
| Triton X-100 | 1 | 0.04 | 0.04 | 0.04 |

### On bead reconstitution of MsbA in peptidiscs

Reconstitution was performed as previously idescribed ^8^. Briefly, 1 mg/mL MsbA in 20 mM Tris/HCl pH 8, 100 mM NaCl, 0.25 mM DTT 0.15 % UDM was incubated with 250 µl Ni-NTA beads for 1 h. The flow through was collected using a spin column. 1 mg/mL of the peptide was added to the beads and incubated for 30 min. Afterwards the beads were washed with 2 column volumes (CV) of Wash Buffer (20 mM Tris/ HCl pH 8, 100 mM NaCl) and the reconstituted MsbA was eluted with 2 CV Elution Buffer (20 mM Tris/ HCl pH 8, 100 mM NaCl, 250 mM Imidazole pH 8).

### Expression and purification of membrane scaffold proteins

pMSP1D1 (Addgene plasmid #20061; http://n2t.net/addgene:20061; RRID:Addgene_20061), pMSP1E3D1 (Addgene plasmid #20066; http://n2t.net/addgene:20066; RRID:Addgene_20066) and pMSP2N2 (Addgene plasmid #29520; http://n2t.net/addgene:29520; RRID:Addgene_29520) plasmids were a gift from the Sligar lab ^29,30,40,41^. They were transformed in BL21(DE3) competent *E. coli* cells and purified as described. Briefly, freshly transformed cells were grown in Terrific Broth (TB) medium supplemented with 10 µg/mL kanamycin and antifoam at 37 °C to an OD_600_ of 0.4 to 0.6. Expression was induced by adding 1 mM IPTG. Cells were harvested after 4 hours, resuspended in 20 mM phosphate buffer pH 7.4 containing 1 mM PMSF, approx. 5 mg DNAse I and 1 % Triton, and lysed via sonication (25 % amplitude for 5 minutes, 25 seconds on, 5 seconds off). Cell debris was removed via centrifugation for 30 minutes at 30.000 g at 4 °C. The lysate was filtered (0.2 µm pore size) and loaded on a HisTRAP^TM^ HP (1 mL) column equilibrated with 40 mM Tris/HCl pH 7.4, 300 mM NaCl. The column was washed stepwise with 10 CV of buffer A (40 mM Tris/HCl pH 8, 300 mM NaCl, 1% Triton), buffer B (40 mM Tris/HCl pH 8, 300 mM NaCl, 50 mM Na-cholate, 20 mM imidazole), buffer C (40 mM Tris/HCl, 300 mM NaCl, 50 mM imidazole) and finally eluted with 5 CVs of elution buffer (40 mM Tris/HCl pH 8, 300 mM NaCl, 400 mM imidazole). The sample was dialyzed overnight at 4°C against dialysis buffer (20 mM Tris/HCl pH 7.4, 100 mM NaCl, 0.5 mM EDTA), concentrated to 15 mg/mL, using a 10 kDa MWCO Amicon concentrator, flash frozen and stored at −80 °C.

### Reconstitution of MsbA in nanodiscs

E. *coli* polar lipids (EPL, Avanti E.*coli* Extract Polar, 100600C) were dried and the lipid film was resuspended in 25 mM Tris/HCl pH 8, 150 mM NaCl, 100 mM Na-cholate to a final concentration of 50 mM. For the EPL:PC MSP1D1 Nanodiscs EPL was mixed with DSPC (Avanti) in a 1:1 molar ratio. Purified MsbA in DDM was reconstituted into nanodiscs using a molar ratio of 1:7.5:200 of MsbA:MSP1D1/MSP1E3D1:lipids and a molar ratio of 1:7.5:750 of MsbA:MSP2N2:lipids. The final Na-cholate concentration was adjusted to 25 mM and the reconstitution mixture was incubated for 1 hour at 4°C while rocking. Equilibrated Bio-Beads SM-2 (65% (w/v)) were added, and the mixture was incubated overnight at 4 °C with gentle agitation. The sample was collected from the Bio-Beads and loaded nanodiscs were separated from empty nanodiscs, MSP and aggregates via size exclusion chromatography, using Superdex 200 Increase 10/300 GL column, equilibrated in SEC buffer (25 mM Tris/HCl pH 8, 150 mM NaCl). Reconstituted nanodiscs were concentrated to 1.5 – 2.5 mg/mL, using a 100 kDa MWCO Amicon concentrator and were either directly used for cryo-EM and ATPase activity assays or flash frozen, and stored at −80 °C.

### A8-35 reconstitution after detergent solubilization

10 mg/mL of MsbA, purified in DDM, was mixed with Amphipol A8-35 in 1:3 molar ratio (protein:amphipol). The mixture was incubated for 3 h at 4 °C while gently stirring. Afterwards, the mixture was added to Bio-Beads (SM2-Adsorbent) and incubated overnight at 4 °C while stirring. The sample was further purified via size exclusion chromatography, using Superdex 200 Increase 5/150 GL column, and the peak fractions were concentrated to 10-14 mg/mL for cryo-EM.

### Direct extraction from the membrane with A18

The membrane fraction (obtained as described in the detergent section) was resuspended in 3 % A18 solution and solubilized overnight at 4 °C while gently stirring. The protein was further purified as explained in the detergent purification section, just using detergent free buffers. After size exclusion chromatography, the peak fractions were pooled and concentrated to 1.8-2.5 mg/mL for cryo-EM.

### ATPase activity measurement

The activity assay was performed as described ^17^. The protein reaction contained 1 μg of purified protein in detergent or 0.2 μg MsbA in lipid nanodiscs in the corresponding SEC buffer. The reaction was started by adding 2 mM MgATP (2 mM ATP, 4 mM MgCl_2_) and incubated for 30 min at 37 °C. The reaction was stopped by adding 50 μL of 12 % (w/v) SDS to 50 μL reaction. To start the colour development, 100 μL of a solution containing equal volumes of 12 % (w/v) ascorbic acid in 1 M HCl and 2 % (w/v) ammonium molybdate in 1 M HCl was added and incubated for 5 min at room temperature. To complete the detection reaction, 150 μL of a solution containing 25 mM sodium citrate, 2 % (w/v) sodium metaarsenite, and 2 % (v/v) acetic acid was added and incubated for 20 min at room temperature. Absorbance at 850 nm was measured in a SpectraMax M5 spectrophotometer (Molecular Devices). Potassium phosphate (0.05 mM to 0.6 mM) was used as a standard for determining the concentration of released phosphate.

### Cryo-EM sample preparation and data acquisition

Cryo-EM grid preparation was done as previously described ^3^. Briefly, the samples were applied on glow discharged (15 mA, 45 sec) C-flat holey carbon grids at concentrations listed in Table 2 and blotted for 6-10s at blot-force 20, using the Vitrobot Mark 4 (Thermo Fisher) at a relative humidity of 100% and 4 °C. Images were collected on a 200 kV Glacios transmission electron microscope (Thermo Fisher), equipped with Falcon 4 direct electron detector and Selectris energy filter, at 165k magnification (pixel size 0.68 Å) with a slit size of 10 eV. The total dose was set to 50 e-/Å^2^. Movies were collected automatically with EPU in AFIS mode (aberration-free image shift).

**Table 2:**
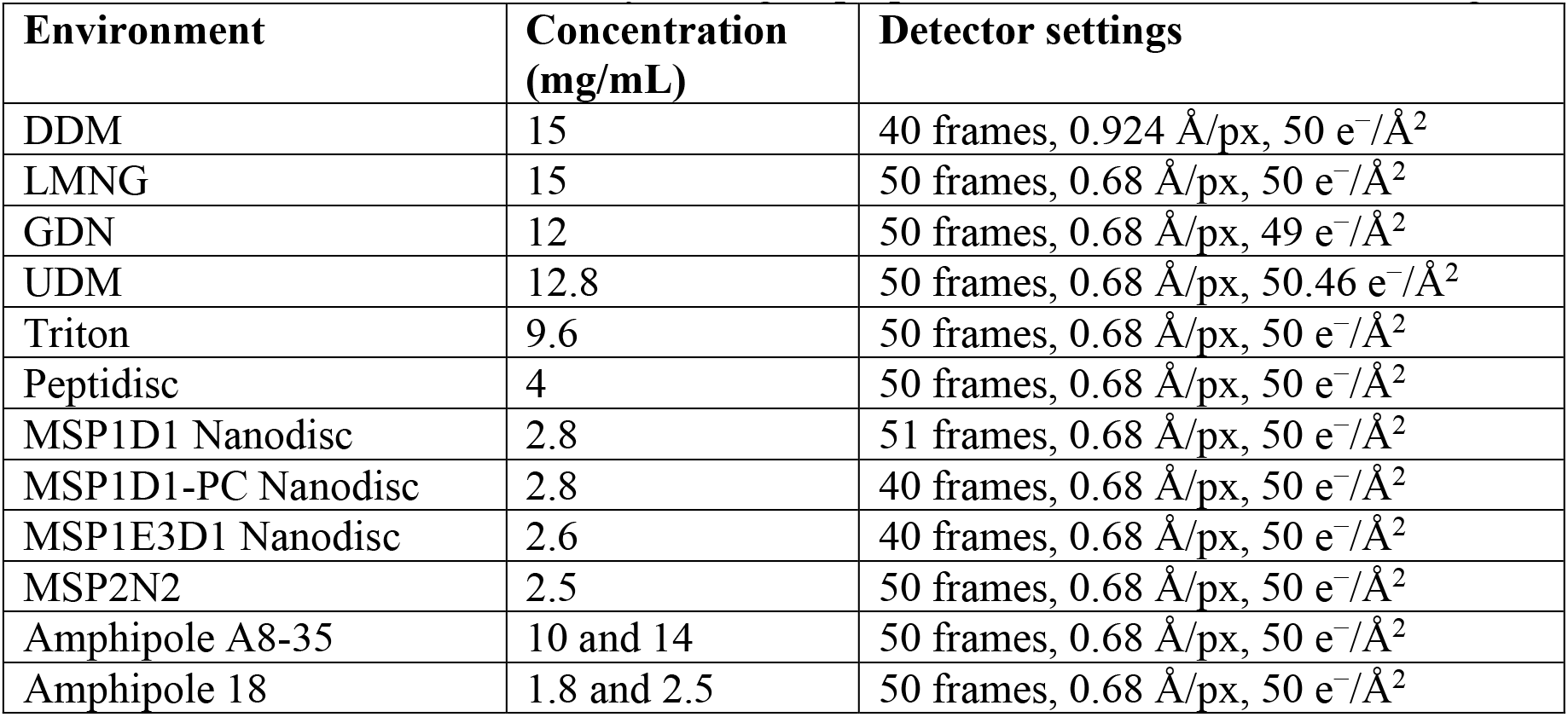
Protein concentration for cryo-EM grid preparation and data collection settings.

| Environment | Concentration (mg/mL) | Detector settings |
| --- | --- | --- |
| DDM | 15 | 40 frames, 0.924 Å/px, 50 e<sup>-</sup>/Å<sup>2</sup> |
| LMNG | 15 | 50 frames, 0.68 Å/px, 50 e<sup>-</sup>/Å<sup>2</sup> |
| GDN | 12 | 50 frames, 0.68 Å/px, 49 e<sup>-</sup>/Å<sup>2</sup> |
| UDM | 12.8 | 50 frames, 0.68 Å/px, 50.46 e<sup>-</sup>/Å<sup>2</sup> |
| Triton | 9.6 | 50 frames, 0.68 Å/px, 50 e<sup>-</sup>/Å<sup>2</sup> |
| Peptidisc | 4 | 50 frames, 0.68 Å/px, 50 e<sup>-</sup>/Å<sup>2</sup> |
| MSP1D1 Nanodisc | 2.8 | 51 frames, 0.68 Å/px, 50 e<sup>-</sup>/Å<sup>2</sup> |
| MSP1D1-PC Nanodisc | 2.8 | 40 frames, 0.68 Å/px, 50 e<sup>-</sup>/Å<sup>2</sup> |
| MSP1E3D1 Nanodisc | 2.6 | 40 frames, 0.68 Å/px, 50 e<sup>-</sup>/Å<sup>2</sup> |
| MSP2N2 | 2.5 | 50 frames, 0.68 Å/px, 50 e<sup>-</sup>/Å<sup>2</sup> |
| Amphipole A8-35 | 10 and 14 | 50 frames, 0.68 Å/px, 50 e<sup>-</sup>/Å<sup>2</sup> |
| Amphipole 18 | 1.8 and 2.5 | 50 frames, 0.68 Å/px, 50 e<sup>-</sup>/Å<sup>2</sup> |

### Data processing

All movies were recorded at a calibrated pixel size of 0.68 Å/px in EER format, except the initial DDM dataset, which was collected at 0.924 Å/px. The respective dose for each dataset is given in Table 2. All processing steps were performed with CryoSPARC (v.4) ^26^, starting with patch-based motion correction ^42^ and patch-based CTF estimation during live preprocessing. Micrographs were curated, using a CTF cutoff at 5 Å. The remaining micrographs were subjected to a sequential picking procedure, by combining particle positions from Blob Picker, Template Picker and Topaz Picker ^43^. Duplicate particles were removed, using the default 20 Å distance parameter. Picked particles were extracted with a box size of 384 px and binned 4 times to 96 px for generating initial 2D classes. After three consecutive rounds of 2D classification, the particle stack was used to create ab initio reconstructions with three to six classes. Heterogeneous refinement (HR) and non-uniform refinement (NUR) ^44^, were used to separately refine the different MsbA conformers. Well-aligning reconstructions were re-extracted without binning and subjected to further rounds of HR and NUR using C2 symmetry. A final round of local refinement with C2 symmetry was additionally applied where needed. A representative processing scheme is given in Supplementary Fig. 7. Details on per-dataset collection settings are provided in Table 2.

### Model building and refinement

For the narrow MsbA conformations, PDB:7PH2 ^18^ was used as a starting model. The wide MsbA conformations were automatically built in ModelAngelo ^45^. The models were rigid-body fitted in each map in UCSF Chimera ^46^ and manually inspected and rebuilt in Coot (v.0.9) ^47^. Refinement was performed using *phenix.real_space_refine* within Phenix (v.1.19) ^48^.

Validation reports were automatically generated within Phenix using MolProbity ^49^. All density maps and models were deposited in the Electron Microscopy Data Bank (EMDB) and the Protein Data Bank (PDB). The PDB IDs are listed in Tab. S1, alongside with the statistics on model building and refinement.

## Supplementary Information

**Supplementary Table 1.** Cryo-EM data collection and processing statistics.

|  | **MsbA-DDM wide** | **MsbA-LMNG narrow** | **MsbA- LMNG wide** | **MsbA-GDN narrow** | **MsbA-GDN wide** |
| --- | --- | --- | --- | --- | --- |
| **Data Collection** |  |  |  |  |  |
| Accession number | EMD-XXXXX | EMD-XXXX | EMD-XXXX | EMD-XXX |  |
| Magnification | 165,000 | 165,000 | 165,000 | 165,000 | 165,000 |
| Voltage / kV | 200 | 200 | 200 | 200 | 200 |
| Dose / e-Å<sup>-2</sup> | 50 | 50 | 50 | 50 | 50 |
| Pixel size / Å | 0.68 | 0.68 | 0.68 | 0.68 | 0.68 |
| Defocus range / μm | -2.0 to -0.8 | -2.0 to -0.8 | -2.0 to -0.8 | -2.0 to -0.8 | -2.0 to -0.8 |
| Recorded movies | 4642 | 7919 | 7919 | 4542 | 4542 |
| Final particle images | 423945 | 285784 | 383680 | 132006 | 206364 |
| Microscope | Glacios | Glacios | Glacios | Glacios | Glacios |
| Camera | Falcon 4 | Falcon 4 | Falcon 4 | Falcon 4 | Falcon 4 |
| Energy Filter | Selectris | Selectris | Selectris | Selectris | Selectris |
| **Image Processing** |  |  |  |  |  |
| Initial model | ModelAngelo | 7PH2 | ModelAngelo | 7PH2 | ModelAngelo |
| Processing software | cryoSPARC (v.4) | cryoSPARC (v.4) | cryoSPARC (v.4) | cryoSPARC (v.4) | cryoSPARC (v.4) |
| Symmetry imposed | C2 | C2 | C2 | C2 | C2 |
| Resolution Å | 3.6 | 3.8 | 3.8 | 3.8 | 3.6 |
| Applied B-factor / Å<sup>2</sup> | 175.6 | 174.1 | 147.8 | 145.5 | 140.8 |

| **Model Refinement** |  |  |  |  |  |
| --- | --- | --- | --- | --- | --- |
| PDB accession | XXXX | XXXX | XXXX | XXXX | XXXX |
| Validation |  |  |  |  |  |
| FSCmap-to-model(0.143) / Å | X.X |  |  |  |  |
| MolProbity score | 1.86 | 1.66 | 1.76 | 2.17 | 1.88 |
| Clash Score | 6.44 | 6.32 | 5.64 | 16.15 | 9.27 |
| Composition |  |  |  |  |  |
| Atoms | 8924 | 8865 | 8822 | 8865 | 8822 |
| Protein residues | 1138 | 1126 | 1138 | 1126 | 1138 |
| Ligands | 3PE: 2 | JSG: 1 | - | JSG: 1 | - |
| Bonds (R.M.S.D.) |  |  |  |  |  |
| Length (Å) | 0.005 | 0.004 | 0.004 | 0.005 | 0.004 |
| Angles (°) | 0.843 | 0.706 | 0.738 | 1.001 | 0.685 |
| B-factors (min/max/mean) |  |  |  |  |  |
| Protein residues | 32.28/199.61/101.02 | 74.06/167.44/118.97 | 23.70/140.24/74.48 | 38.41/161.96/96.37 | 17.74/132.22/65.41 |
| Ligand | 20.00/20.00/20.00 | 30.00/30.00/30.00 | - | 30.00/30.00/30.00 | - |
| Ramachandran plot (%) |  |  |  |  |  |
| Favored | 91.53 | 95.45 | 92.77 | 92.69 | 94.27 |
| Allowed | 8.38 | 4.46 | 7.14 | 7.22 | 5.73 |
| Outliers | 0.09 | 0.09 | 0.09 | 0.09 | 0.00 |
| Rotamer outliers (%) | 0.31 | 0.52 | 0.10 | 0.31 | 0.62 |

|  | **MsbA-Triton wide** | **MsbA-Triton wide+** | **MsbA- UDM wide** | **MsbA-UDM wide+** | **MsbA-MSP1D1 narrow** |
| --- | --- | --- | --- | --- | --- |
| **Data Collection** |  |  |  |  |  |
| Accession number | EMD-XXXXXX | EMD-XXXX | EMD-XXXX | EMD-XXX |  |
| Magnification | 165,000 | 165,000 | 165,000 | 165,000 | 165,000 |
| Voltage / kV | 200 | 200 | 200 | 200 | 200 |
| Dose / e-Å<sup>-2</sup> | 50 | 50 | 50 | 50 | 50 |
| Pixel size / Å | 0.68 | 0.68 | 0.68 | 0.68 | 0.68 |
| Defocus range / μm | -2.0 to -0.8 | -2.0 to -0.8 | -2.0 to -0.8 | -2.0 to -0.8 | -2.0 to -0.8 |
| Recorded movies | 5094 | 5094 | 11655 | 11655 | 8819 |
| Final particle images | 92493 | 96119 | 210730 | 162890 | 163040 |
| Microscope | Glacios | Glacios | Glacios | Glacios | Glacios |
| Camera | Falcon 4 | Falcon 4 | Falcon 4 | Falcon 4 | Falcon 4 |
| Energy Filter | Selectris | Selectris | Selectris | Selectris | Selectris |
| **Image Processing** |  |  |  |  |  |
| Initial model | ModelAngelo | ModelAngelo | ModelAngelo | ModelAngelo | 7PH2 |
| Processing software | cryoSPARC (v.4) | cryoSPARC (v.4) | cryoSPARC (v.4) | cryoSPARC (v.4) | cryoSPARC (v.4) |
| Symmetry imposed | C2 | C2 | C2 | C2 |  |
| Resolution Å | 4.0 | 3.8 | 4.1 | 4.2 | 3.9 |
| Applied B-factor / Å<sup>2</sup> | 159.1 | 135.9 | 240.9 | 259.3 | 162.2 |
| **Model Refinement** |  |  |  |  |  |
| PDB accession | XXXX | XXXX | XXXX | XXXX | XXXX |
| Validation |  |  |  |  |  |
| FSCmap-to-model(0.143) / Å | X.X |  |  |  |  |
| MolProbity score | 2.04 | 2.05 | 2.32 | 1.78 | 1.77 |
| Clash Score | 14.86 | 13.97 | 23.91 | 8.32 | 7.49 |
| Composition |  |  |  |  |  |
| Atoms | 8822 | 8822 | 8822 | 8822 | 8865 |
| Protein residues | 1138 | 1138 | 1138 | 1138 | 1126 |
| Ligands | - | - | - | - | JSG: 1 |
| Bonds (R.M.S.D.) |  |  |  |  |  |
| Length (Å) | 0.004 | 0.004 | 0.007 | 0.005 | 0.006 |
| Angles (°) | 0.757 | 0.729 | 0.785 | 0.989 | 1.264 |
| B-factors (min/max/mean) |  |  |  |  |  |
| Protein residues | 108.66/222.40/157.18 | 69.78/182.02/116.80 | 163.96/306.89/222.53 | 143.50/287.03/202.50 | 65.28/195.17/133.52 |
| Ligand | - | - | - | - | 30.00/30.00/30.00 |
| Ramachandran plot (%) |  |  |  |  |  |
| Favored | 94.71 | 94.18 | 93.30 | 95.24 | 94.83 |
| Allowed | 5.03 | 5.73 | 6.53 | 4.50 | 5.17 |
| Outliers | 0.26 | 0.09 | 0.18 | 0.26 | 0.00 |
| Rotamer outliers (%) | 0.31 | 0.31 | 1.03 | 0.52 | 0.63 |

|  | **MsbA-MSP1D1-**<br>**PC narrow** | **MsbA-**<br>**MSP1E3D1**<br>**narrow** | **MsbA- MSP2N2**<br>**narrow** | **MsbA-MSP2N2**<br>**wide** | **MsbA-peptidisc**<br>**narrow-** |
| --- | --- | --- | --- | --- | --- |
| **Data Collection** |  |  |  |  |  |
| Accession number | EMD-XXXXX | EMD-XXXX | EMD-XXXX | EMD-XXX |  |
| Magnification | 165,000 | 165,000 | 165,000 | 165,000 | 165,000 |
| Voltage / kV | 200 | 200 | 200 | 200 | 200 |
| Dose / e-Å<sup>-2</sup> | 50 | 50 | 50 | 50 | 50 |
| Pixel size / Å | 0.68 | 0.68 | 0.68 | 0.68 | 0.68 |
| Defocus range /<br>µm | -2.0 to -0.8 | -2.0 to -0.8 | -2.0 to -0.8 | -2.0 to -0.8 | -2.0 to -0.8 |
| Recorded movies | 4866 | 6430 | 66501 | 66501 | 17117 |
| Final particle<br>images | 89100 | 304832 | 346172 | 140572 | 188903 |
| Microscope | Glacios | Glacios | Glacios | Glacios | Glacios |
| Camera | Falcon 4 | Falcon 4 | Falcon 4 | Falcon 4 | Falcon 4 |
| Energy Filter | Selectris | Selectris | Selectris | Selectris | Selectris |
| **Image Processing** |  |  |  |  |  |
| Initial model | 7PH2 | 7PH2 | 7PH2 | ModelAngelo | 7PH2 |
| Processing<br>software | cryoSPARC (v.4) | cryoSPARC (v.4) | cryoSPARC (v.4) | cryoSPARC (v.4) | cryoSPARC (v.4) |
| Symmetry<br>imposed | C2 | C2 | C2 | C2 | C2 |
| Resolution Å | 3.8 | 3.3 | 3.9 | 4.4 | 3.8 |
| Applied B-factor /<br>Å<sup>2</sup> | 156.1 | 145.2 | 156.6 | 232.9 | 161.4 |
| **Model**<br>**Refinement** |  |  |  |  |  |
| PDB accession | XXXX | XXXX | XXXX | XXXX | XXXX |
| Validation |  |  |  |  |  |
| FSCmap-to-model(0.143) / Å | X.X |  |  |  |  |
| MolProbity score | 1.86 | 1.76 | 1.70 | 1.81 | 1.96 |
| Clash Score | 9.51 | 7.27 | 5.55 | 6.87 | 10.60 |
| Composition |  |  |  |  |  |
| Atoms | 8799 | 8966 | 8864 | 8822 | 8865 |
| Protein residues | 1118 | 1126 | 1126 | 1138 | 1126 |
| Ligands | JSG: 1 | JSG: 1, 3PE: 2 | JSG: 1 | - | JSG: 1 |
| Bonds (R.M.S.D.) |  |  |  |  |  |
| Length (Å) | 0.006 | 0.007 | 0.005 | 0.005 | 0.006 |
| Angles (°) | 1.108 | 1.267 | 1.097 | 1.039 | 1.144 |
| B-factors (min/max/mean) |  |  |  |  |  |
| Protein residues | 83.49/194.47/134.86 | 0.24/127.62/56.62 | 72.52/207.71/133.82 | 157.88/322.58/215.64 | 93.82/204.31/146.79 |
| Ligand | 30.00/30.00/30.00 | 28.70/33.27/30.39 | 117.66/117.66/117.66 | - | 30.00/30.00/30.00 |
| Ramachandran plot (%) |  |  |  |  |  |
| Favored | 94.79 | 94.83 | 94.10 | 93.30 | 93.67 |
| Allowed | 5.03 | 5.08 | 5.81 | 6.44 | 6.15 |
| Outliers | 0.18 | 0.09 | 0.09 | 0.26 | 0.18 |
| Rotamer outliers (%) | 0.00 | 0.73 | 0.10 | 0.31 | 0.84 |

|  | **MsbA-peptidisc**<br>**narrow** | **MsbA-A8-35 wide** | **MsbA- A8-35 wide+** | **MsbA-**<br>**Ampipole-18**<br>**wide** | **MsbA-**<br>**Amphipole-18**<br>**wide+** |
| --- | --- | --- | --- | --- | --- |
| **Data Collection** |  |  |  |  |  |
| Accession number | EMD-XXXXX | EMD-XXXX | EMD-XXXX | EMD-XXX |  |
| Magnification | 165,000 | 165,000 | 165,000 | 165,000 | 165,000 |
| Voltage / kV | 200 | 200 | 200 | 200 | 200 |
| Dose / e-Å<sup>-2</sup> | 50 | 50 | 50 | 50 | 50 |
| Pixel size / Å | 0.68 | 0.68 | 0.68 | 0.68 | 0.68 |
| Defocus range /<br>μm | -2.0 to -0.8 | -2.0 to -0.8 | -2.0 to -0.8 | -2.0 to -0.8 | -2.0 to -0.8 |
| Recorded movies | 17117 | 16884 | 16884 | 18842 | 18842 |
| Final particle<br>images | 171149 | 152504 | 101023 | 378262 | 277192 |
| Microscope | Glacios | Glacios | Glacios | Glacios | Glacios |
| Camera | Falcon 4 | Falcon 4 | Falcon 4 | Falcon 4 | Falcon 4 |
| Energy Filter | Selectris | Selectris | Selectris | Selectris | Selectris |
| **Image Processing** |  |  |  |  |  |
| Initial model | 7PH2 | ModelAngelo | ModelAngelo | ModelAngelo | ModelAngelo |
| Processing<br>software | cryoSPARC (v.4) | cryoSPARC (v.4) | cryoSPARC (v.4) | cryoSPARC<br>(v.4) | cryoSPARC<br>(v.4) |
| Symmetry<br>imposed | C2 | C2 | C2 | C2 | C2 |
| Resolution Å | 3.9 | 3.8 | 4.0 | 3.4 | 3.6 |
| Applied B-factor /<br>Å<sup>2</sup> | 181.3 | 111.0 | 124.5 | 128.4 | 148.3 |
| **Model**<br>**Refinement** |  |  |  |  |  |
| PDB accession | XXXX | XXXX | XXXX | XXXX | XXXX |
| Validation |  |  |  |  |  |
| FSCmap-to-model(0.143) / Å | X.X |  |  |  |  |
| MolProbity score | 1.92 | 1.70 | 1.79 | 1.57 | 1.56 |
| Clash Score | 11.71 | 6.09 | 9.44 | 3.30 | 3.85 |
| Composition |  |  |  |  |  |
| Atoms | 8865 | 8822 | 8822 | 8822 | 8822 |
| Protein residues | 1126 | 1138 | 1138 | 1138 | 1138 |
| Ligands | JSG: 1 | - | - | - | - |
| Bonds (R.M.S.D.) |  |  |  |  |  |
| Length (Å) | 0.006 | 0.005 | 0.005 | 0.005 | 0.005 |
| Angles (°) | 1.079 | 0.889 | 1.275 | 0.877 | 1.117 |
| B-factors (min/max/mean) |  |  |  |  |  |
| Protein residues | 92.56/184.76/137.23 | 139.03/261.55/178.76 | 112.04/291.39/178.31 | 9.30/98.50/42.16 | 9.30/98.50/42.16 |
| Ligand | 30.00/30.00/30.00 | - | - | - | - |
| Ramachandran plot (%) |  |  |  |  |  |
| Favored | 95.10 | 94.71 | 95.77 | 93.03 | 94.36 |
| Allowed | 4.81 | 5.11 | 4.23 | 6.88 | 5.56 |
| Outliers | 0.09 | 0.18 | 0.00 | 0.09 | 0.09 |
| Rotamer outliers (%) | 0.21 | 0.41 | 0.62 | 0.10 | 0.21 |

**Supplementary Table 2.** Distance measurements. The distances are labeled in a gradient from cyan (lowest) to blue (highest) distance. Published structures for comparison are shown at the bottom. Triangles indicate direct extractions and circles reconstitutions.

| ID | Distance (E506) | Distance (D512) |
| --- | --- | --- |
| ○ Peptidisc-IF_narrow- | 22.5 | 31.8 |
| ○ Nanodisc_MSP1D1-PC-IF_narrow | 23.1 | 34.8 |
| ○ Nanodisc_MSP1E3D1-IF_narrow | 23.8 | 37.8 |
| ○ Nanodisc_MSP2N2-IF_narrow | 24.3 | 38.1 |
| ○ Nanodisc_MSP1D1-IF_narrow | 29.1 | 40.4 |
| △ GDN-IF_narrow | 31.5 | 40.8 |
| △ LMNG-IF_narrow | 35.3 | 50.1 |
| ○ Peptidisc-IF_narrow | 35.8 | 49.5 |
| ○ Nanodisc_MSP2N2-IF_wide | 51.7 | 64.1 |
| △ GDN-IF_wide | 57.8 | 67.9 |
| △ UDM-IF_wide | 57.9 | 68.5 |
| △ Triton-IF_wide | 58.3 | 66.9 |
| △ LMNG-IF_wide | 60.5 | 68.5 |
| △ DDM-IF_wide | 60.8 | 70.2 |
| ○ Amphipol_A8-35-IF_wide | 60.9 | 71.5 |
| △ Amphipol_18-IF_wide | 64.6 | 72.0 |
| △ UDM-IF_wide+ | 67.9 | 78.7 |
| △ Triton-IF_wide+ | 68.4 | 77.8 |
| ○ Amphipol_A8-35-IF_wide+ | 70.4 | 80.0 |
| △ Amphipol_18-IF_wide+ | 74.0 | 79.1 |
| ○ 5TV4_nanodisc-IF_narrow | 26.1 | 33.9 |
| ○ 6UZ2_peptidisc-IF_narrow | 28.8 | 44.0 |
| △ 8DMO_DDM-IF_wide | 68.2 | 68.4 |

**Supplementary Fig. 1.**
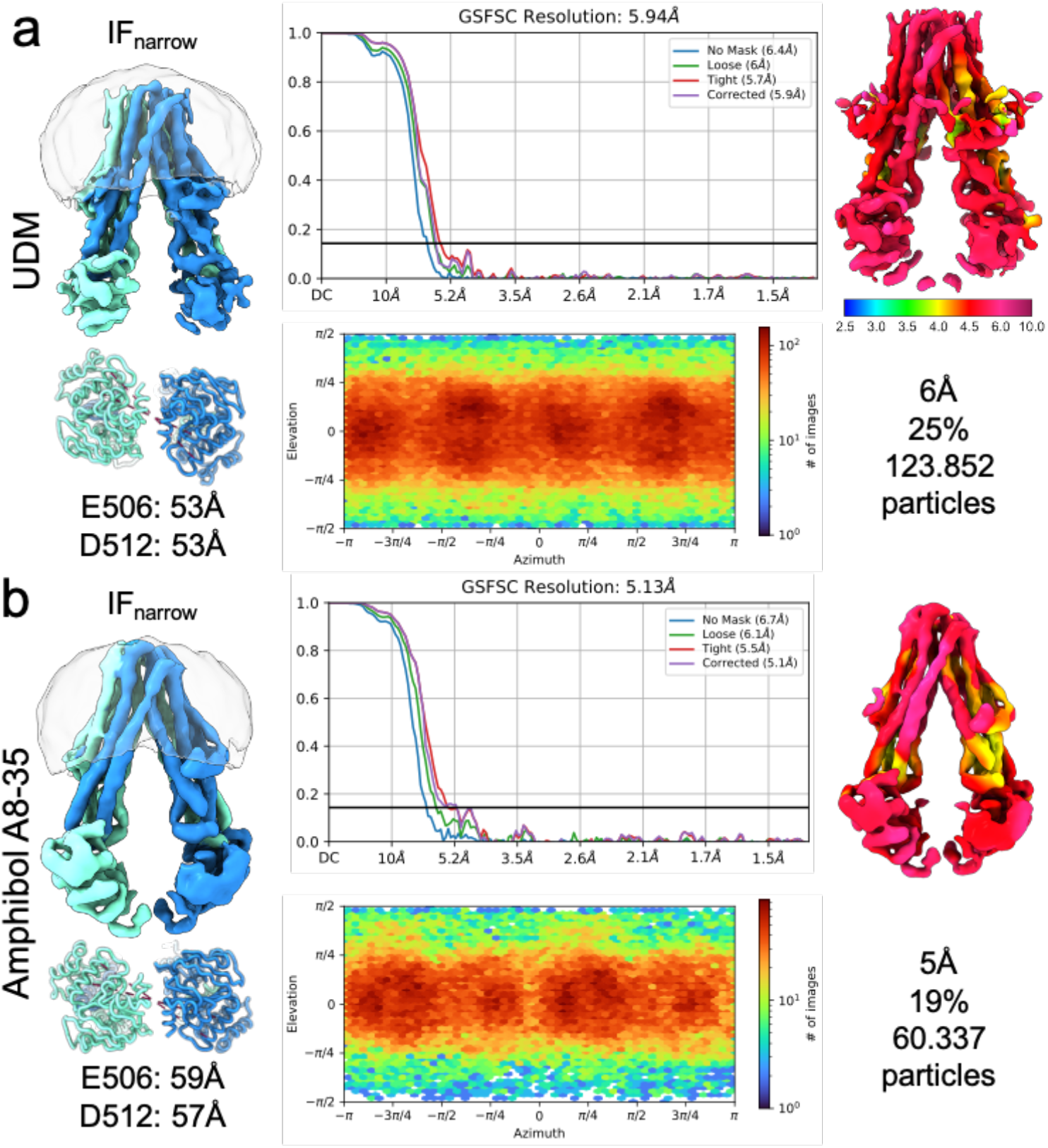
Narrow conformations of MsbA in UDM and amphipol A8-35. Low-resolution density maps for the datasets of UDM and amphipol A8-35. Local resolution representations are visualized on the right side. Particle numbers are presented as values as well as percentages to represent the distribution within the whole data set. The particle orientation as angular distribution plot and FSC (gold-standard, 0.143) are shown for validation.

**Supplementary Fig. 2.**
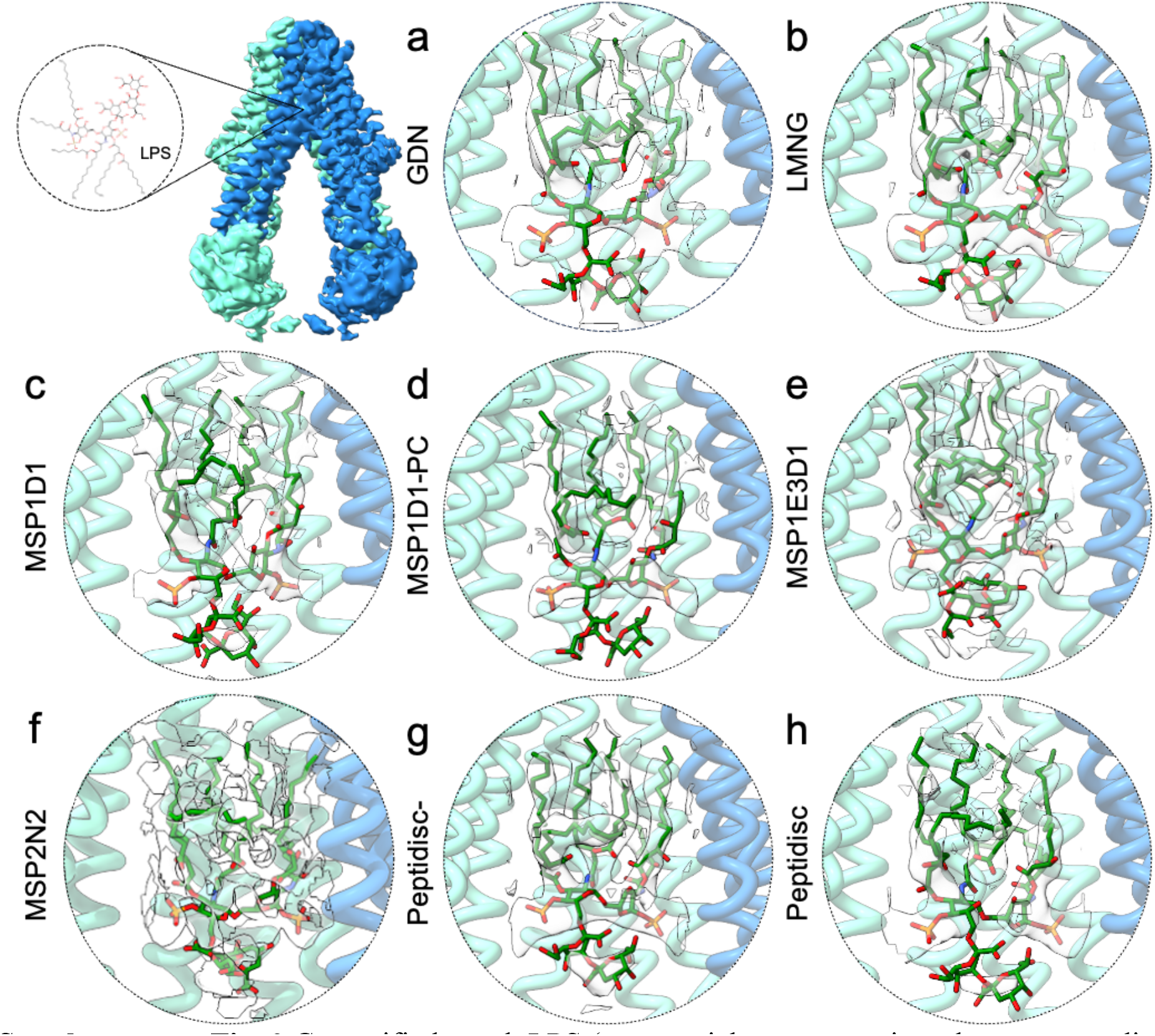
Co-purified rough LPS (green, stick representation; the corresponding semi-transparent density is zoned around the model) is encapsulated within the inner cavity of MsbA, assignable exclusively in its narrow conformations, both in detergent and nanodisc environments. MsbA densities and structures are coloured by chain ID (cyan and blue).

**Supplementary Fig. 3.**
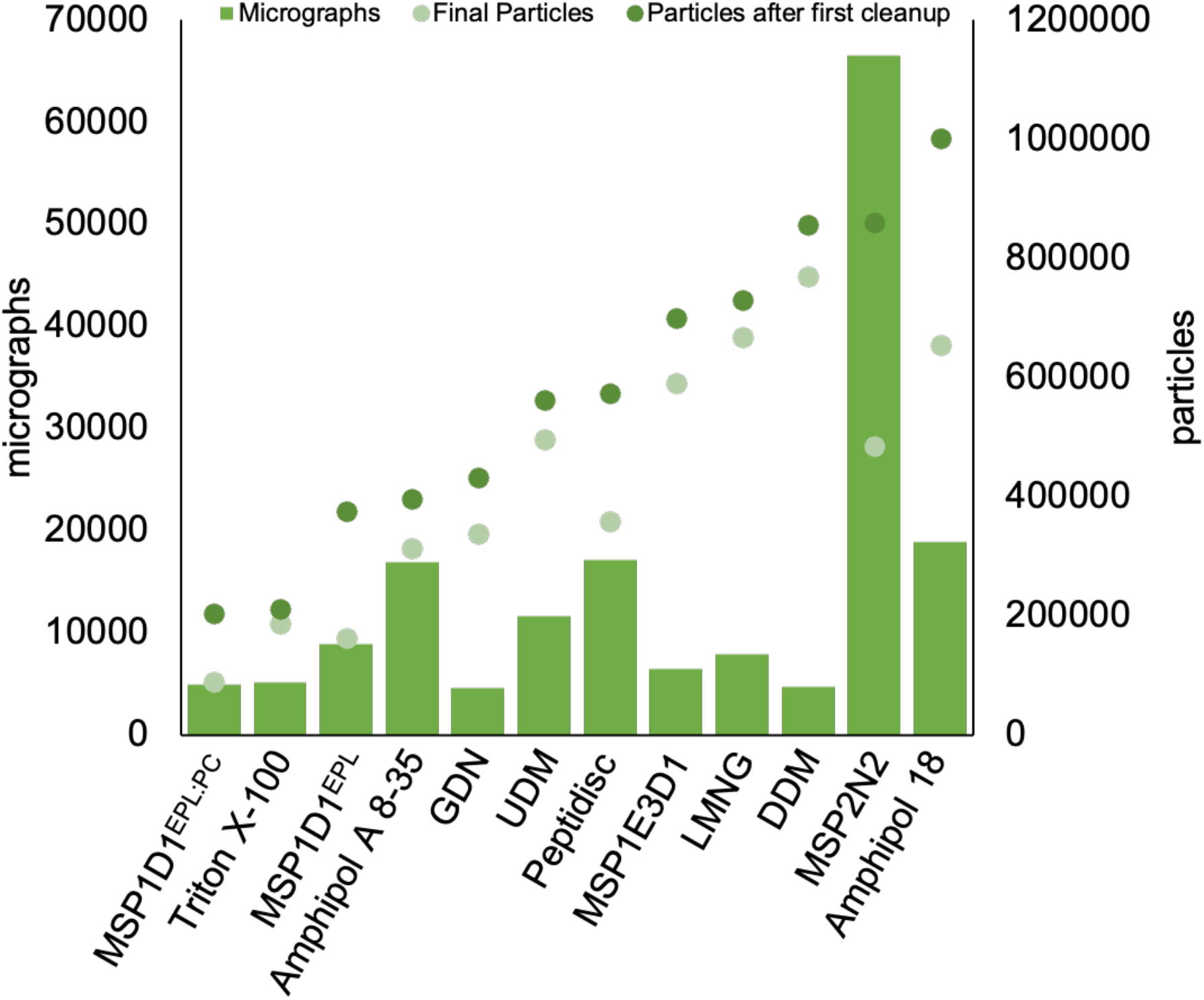
Representation of Particle and Micrograph distribution. Bars indicate the recorded micrographs, dots in dark green represent the particles after the first cleanup (2D classification), and dots in light green – final particles.

**Supplementary Fig. 4.**
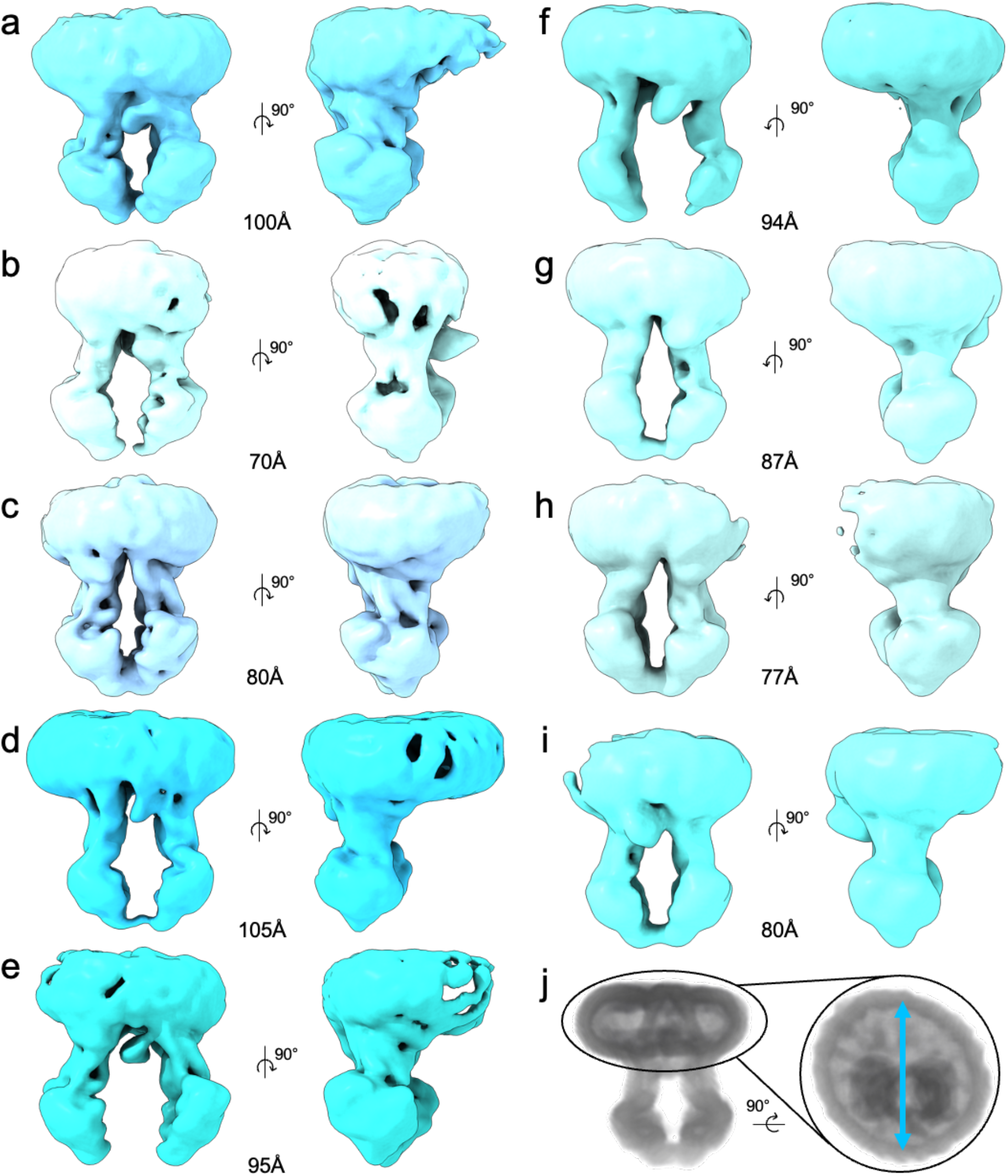
Variation of Nanodisc sizes in the MSP2N2 dataset. (a-i) Surface representation of refined maps of different conformational states as well as Nanodisc sizes of MSP2N2. Front and side view (right or left rotation by 90° of the map indicated with arrows). Approximate measurement of the nanodisc size in Å. (j) Measurement of the Nanodisc size as indicated from the top by 90° rotation. Volume representation of the map to show the location of the protein in the Nanodisc.

**Supplementary Fig. 5.**
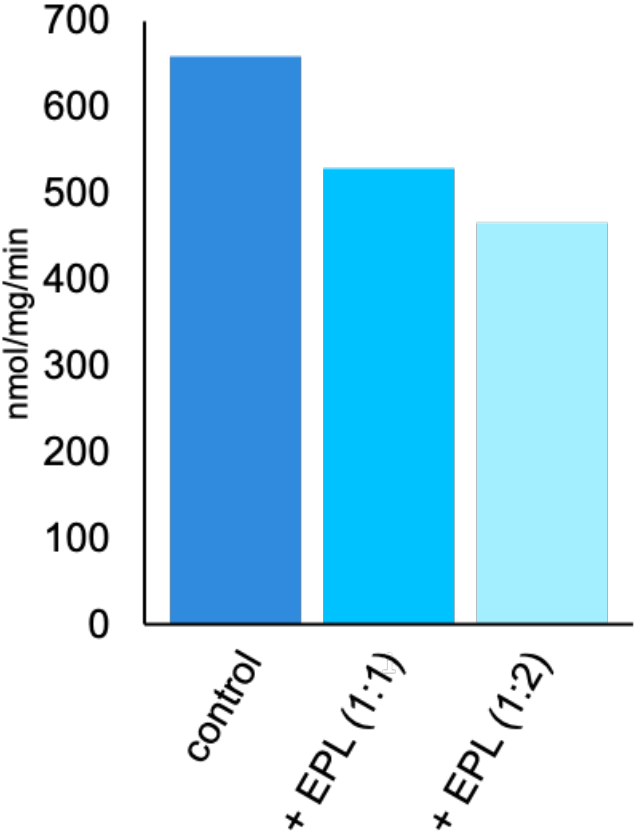
ATPase activity measurement of MsbA in LMNG with added Lipids. As a control the basal activity of MsbA in LMNG without the addition of lipids was measured. To test the influence of Lipids *E. coli* polar lipids were added to MsbA in LMNG in the ratio 1:1 and 1:2(protein:lipis, w/w).

**Supplementary Fig. 6.**
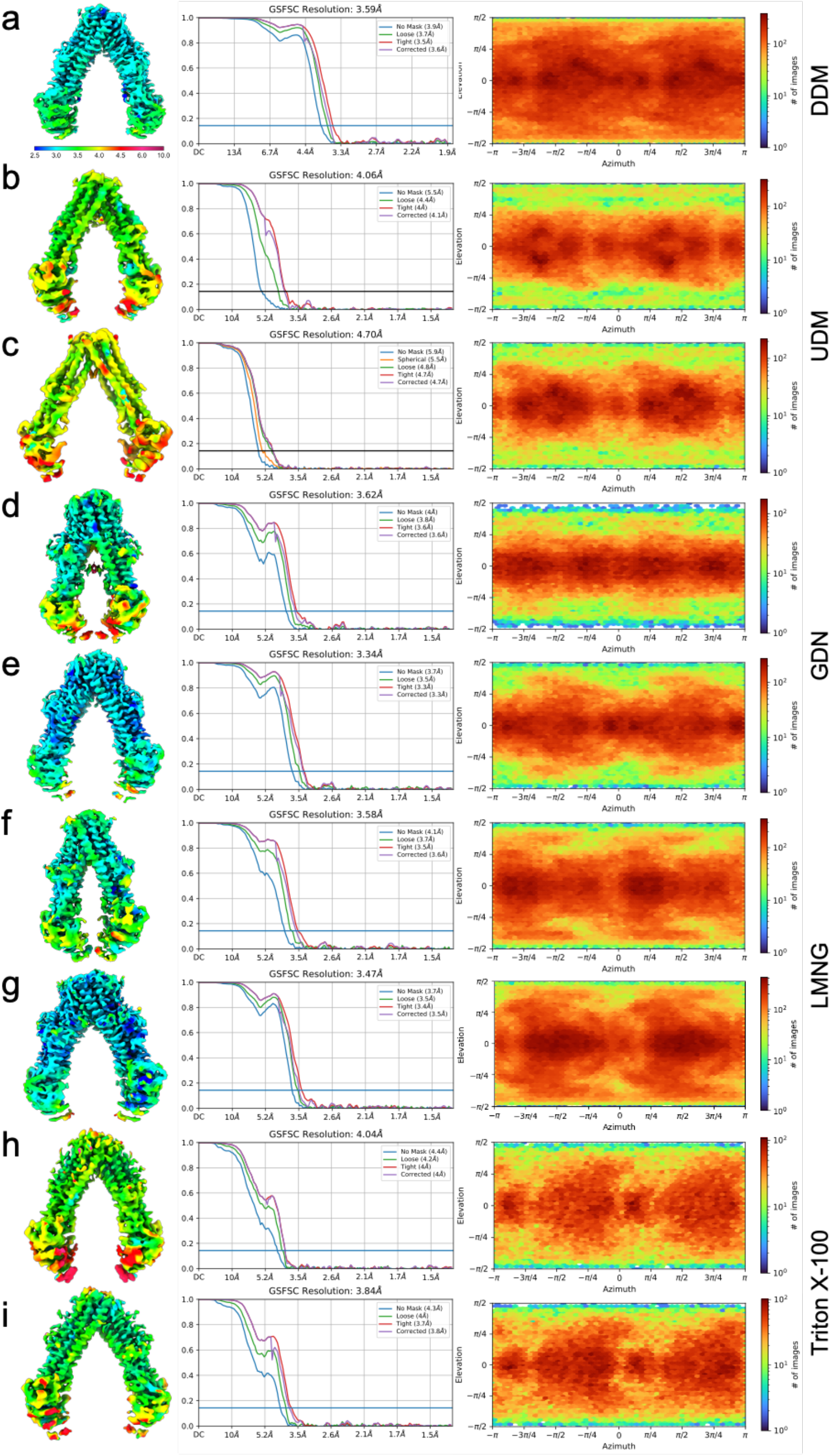
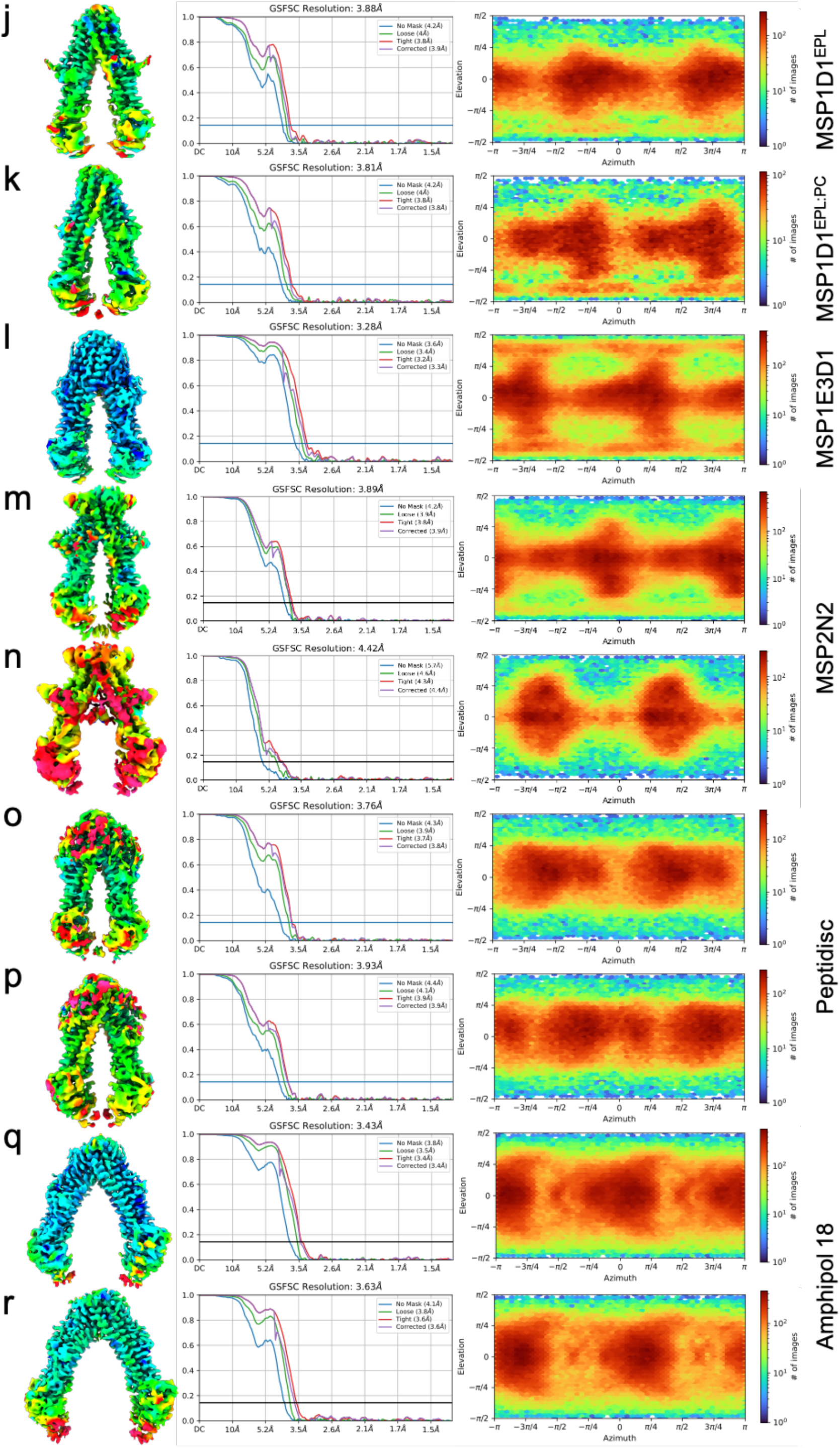
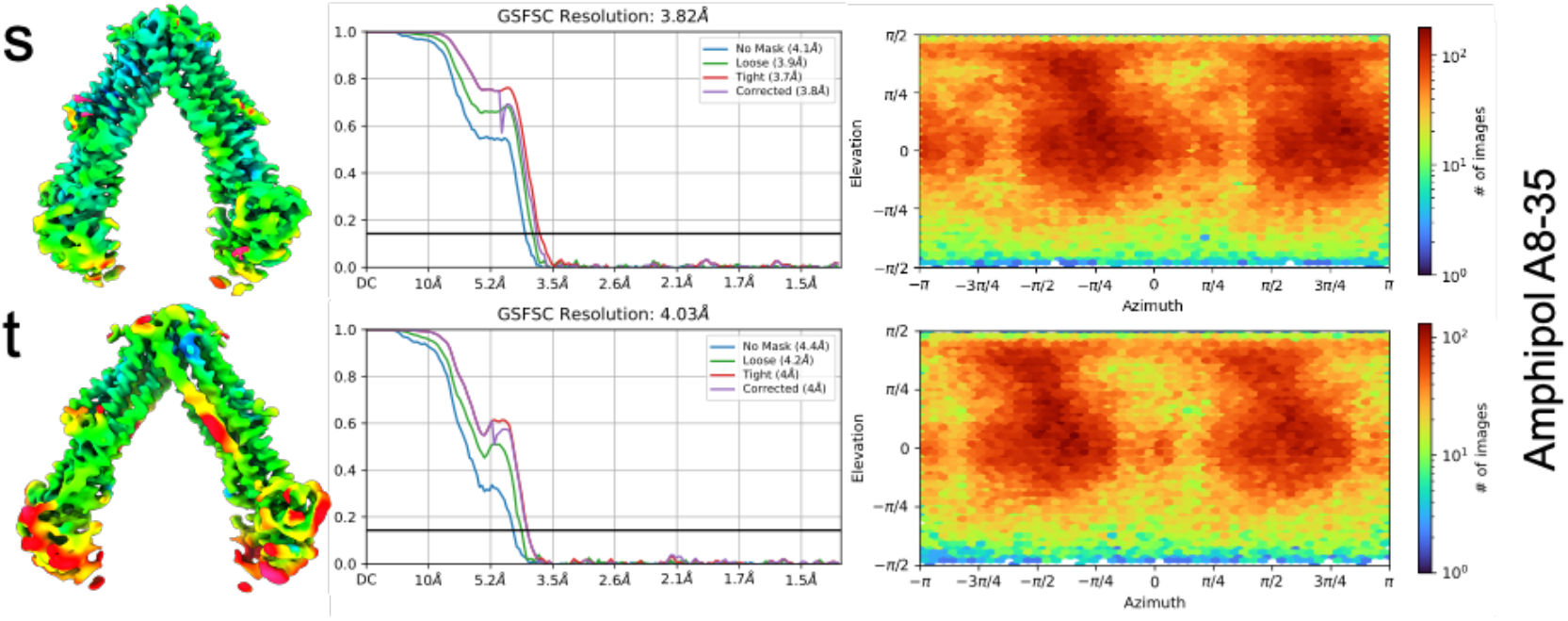
Quality of MsbA maps. Cryo-EM density maps are coloured according to the local-resolution estimation. Gold-standard FSC curves (FSC = 0.143) are displayed together with angular distribution plots for all solved structures.

**Supplementary Fig. 7.**
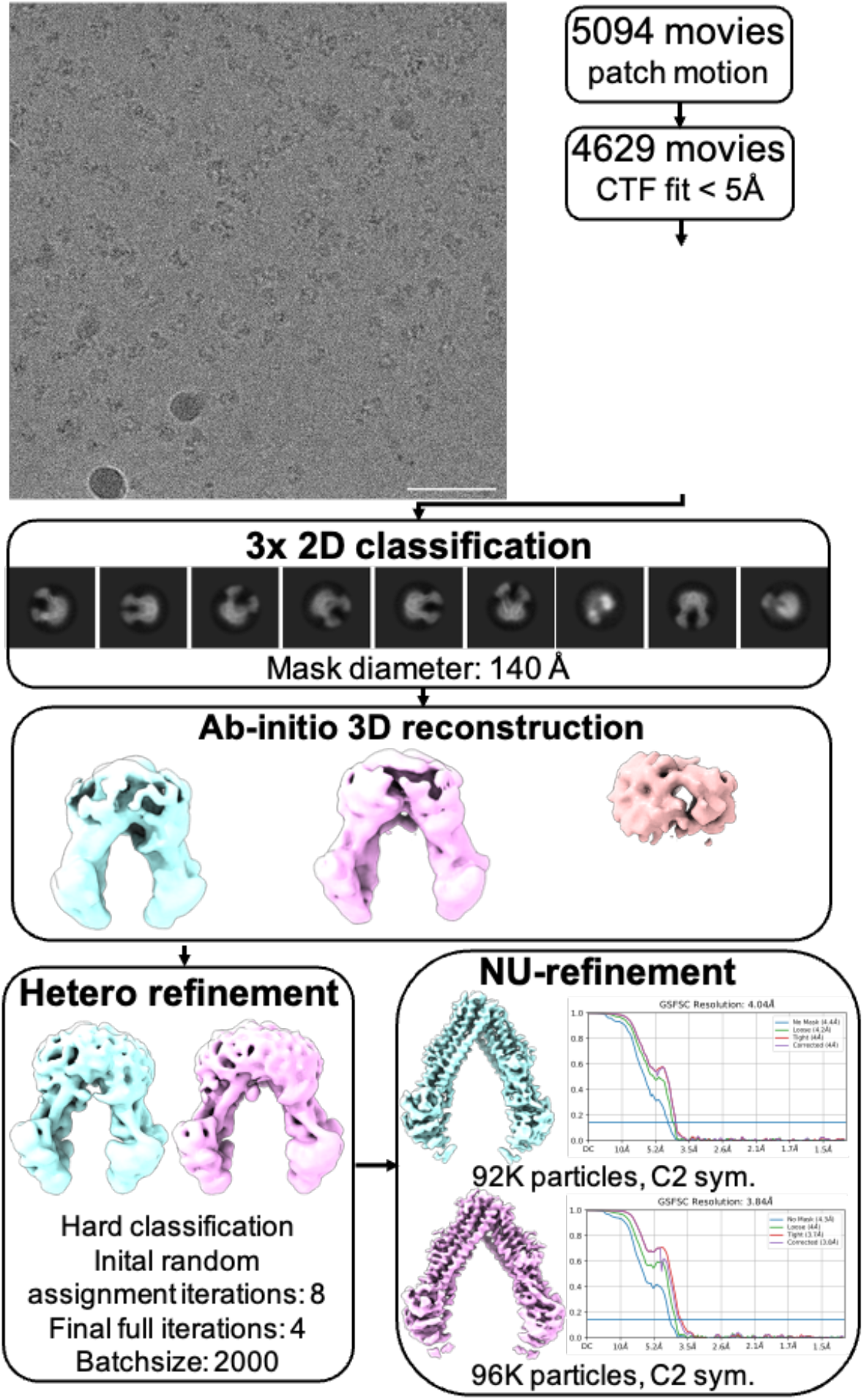
Cryo-EM processing workflow. Representative processing workflow for MsbA in Triton X-100 with cryoSPARC and cryoSPARC live. All datasets were processed in the same manner; only for the lipid nanodisc datasets refinement with custom masking was included.

## Notes

### Competing Interest Statement

The authors have declared no competing interest.

